# Intratumoral amino acid insufficiency limits CD8^+^ T-cell effector function

**DOI:** 10.1101/2025.04.28.651077

**Authors:** Yan-Ting Chen, Ya-Hui Lin, Jahan Rahman, Justin Cross, Natalya N. Pavlova, Santosha A. Vardhana

## Abstract

Loss of effector function is a hallmark of tumor-infiltrating CD8^+^ T-cells that have lost therapeutic efficacy. This impaired capacity occurs despite expression of transcripts encoding cytotoxic proteins, raising the possibility that post-transcriptional suppression of cytotoxic protein synthesis limits anti-tumor immunity. Whether altered protein synthesis contributes to CD8^+^ T-cell dysfunction has not been explored. Here we show that intratumoral amino acid availability restricts the cytotoxic capacity of CD8^+^ TILs by perturbing their ability to sustain protein synthesis. mRNA translation rates in antigen-specific CD8^+^ T-cells were rapidly and specifically suppressed within tumors but not tumor-draining lymph nodes, due to a combination of increased amino acid demand and reduced amino acid availability. Mechanistically, amino acid-dependent uncharging of tRNA^Gln^ in T-cells persistently exposed to antigen was sufficient to suppress protein synthesis in a manner that is independent of either activation of the integrated stress response or suppression of mTORC1 activation. Finally, suppressing intracellular glutaminase activity or ectopically overexpressing the amino acid transporter SLC6A15 was sufficient to restore CD8^+^ T-cell effector function. These results establish a novel mechanism by which nutrient availability in the tumor microenvironment limits T-cell function and demonstrate how enhancing T cell-specific amino acid availability can sustain T-cell effector function and potentiate anti-tumor immunity.

## INTRODUCTION

To engage in serial target cell lysis, cytotoxic CD8^+^ T-cells must synthesize inflammatory cytokines and cytolytic granules at high rates. Nearly sixty years ago, Ingegerd Hellstrom described a paradox in which circulating CD8^+^ T-cells were able to successfully lyse autochthonous tumor cells, but tumor-resident CD8^+^ T-cells were unable to do so^1^. Several molecular drivers of intratumoral T-cell dysfunction have been identified, including the transcriptional and epigenetic programs that drive an ‘exhausted’ T-cell state similar to T-cells from patients with chronic viral infections^2–6^. Notably, tumor-infiltrating dysfunctional CD8^+^ T-cells (TILs) express high levels of transcripts encoding cytotoxic proteins, raising the question of whether effector dysfunction in TILs is regulated at the level of protein synthesis^7^. However, the mechanistic basis for why intratumoral CD8^+^ T-cells fail to sustain the production of effector proteins in response to persistent antigenic challenge has been less well studied.

### mRNA translation is suppressed in antigen-specific CD8^+^ T-cells upon tumor infiltration

Several studies have established common transcriptional, epigenetic, and functional features of exhausted CD8^+^ T-cells isolated from either tumors or mice bearing chronic viral infections, such as the Clone 13 strain of lymphocytic choriomeningitis virus (LCMV-Cl13), including upregulation of inhibitory immunoreceptors and transcription factors as well as a loss of effector cytokine production upon restimulation^8–10^. We confirmed these findings by performing bulk RNA sequencing of CD44^+^CD8^+^ T-cells from mice bearing the Armstrong strain of LCMV (LCMV-Arm), which generates primarily effector CD8^+^ T cells as 7 days post infection (dpi), CD44^+^CD8^+^ T-cells from mice bearing LCMV-Cl13, which generates primarily exhausted T cells at 28 (dpi)^11,12^, or tumor infiltrating CD44^+^CD8^+^ T cells from mice bearing either E0771 breast or B16 melanoma tumors at 21 days following tumor implantation (**Fig. 1a**). Consistent with prior publications^11^, T cells from LCMV-Cl13 mice as well as TILs from B16 and E0771 tumors expressed multiple inhibitory immunoreceptors including *Pdcd1*, *Ctla4*, *Havcr2*, *Lag3*, *Entpd1*, and *Tnfrsf9* as well as known exhaustion-associated transcription factors such as *Nr4a2, Irf8*, and *Tox,* and downregulated memory associated genes including *Tcf7*, *S1pr1*, *S1pr4*, *Sell*, and *Lef1* (**Fig. 1b**). Similarly, both T cells from LCMV-Cl13 and TILs from either B16 or E0771 tumors exhibited loss of cytotoxic protein production upon restimulation **(Extended Data Fig. 1a)**. In contrast to exhausted CD8^+^ T-cells from LCMV-Cl13-infected mice, however, CD8^+^ TILs expressed high levels of transcripts encoding cytotoxic molecules, including *Ifng*, *Tnf, Gzmb*, and *Gzmc,* suggesting that the failure of TILs to sustain effector protein production might be regulated at the level of protein synthesis **(Fig. 1b)**.

**Fig 1.**
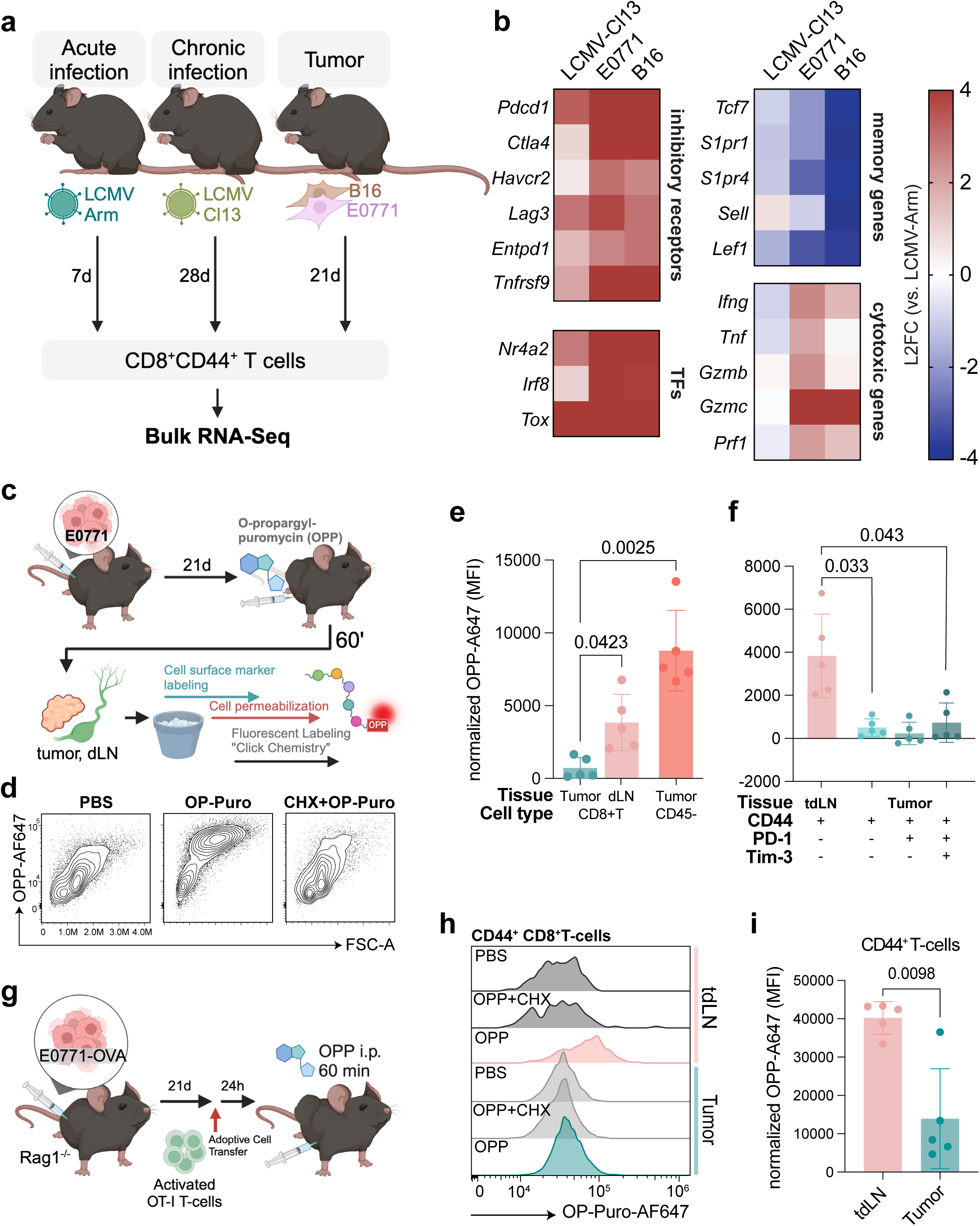
Protein synthesis is suppressed in tumor infiltrating CD8^+^ T-cells *in vivo*. a, Experimental design to profile CD8^+^CD44^+^ T-cell transcriptomics from acute or chronic infection models, as well as in tumors via bulk RNA-seq. **b,** Heatmap showing the expression of cytotoxic cytokines and T cell exhaustion-associated genes in T-cells isolated from LCMV Clone 13-infected mice and intratumoral T-cells from E0771-tumor bearing mice, measured by RNA-seq. Log_2_ Fold Change is calculated relative to the expression in effector T-cells isolated from LCMV Armstrong-infected mice. **c,** Experimental design to measure single cell protein synthesis rates *in vivo*. **d,** OPP-Alexa Fluor 647 (OPP-AF647) signal of CD45^-^ stromal cell population from tumor-bearing mice injected with either PBS, OPP, or CHX pretreatment prior to OPP. **e,** OPP-AF647 Mean fluorescence intensity (MFI) of CD44^+^ T-cells and CD45^-^ stromal cells from the tumor and CD44^+^ T-cells from the tumor-draining lymph node (tdLN), normalized to PBS control. **f,** OPP-AF647 MFI of tumor infiltrating CD44^+^ T-cells in (**e**), stratified by expression of PD-1 and Tim-3 and normalized to PBS control. **g,** Experimental design to measure protein synthesis rates in OT-I TILs 24 hours after adoptive transfer into E0771-LentiLucOS tumor-bearing mice. **h,i,** OPP-AF647 MFI of OT-I TILs described in (**g**), normalized to PBS control and quantified in (**i**). P values were calculated by one-way ANOVA with Dunnett’s multiple comparisons test (**e,f**), or two-tailed Student’s t-test (**i**). Data are presented as the mean ± s.d. of n = 5 independent mice from a representative experiment.

To measure protein synthesis rates in tumor-infiltrating T-cells, we leveraged O-propargyl-puromycin (OPP), an alkyne analog of puromycin that is incorporated into translating polypeptides, providing a single-cell readout of protein synthesis rates when conjugated to a fluorophore by click chemistry^13,14^. We injected OPP intraperitoneally into C57/BL6 mice bearing subcutaneous E0771 tumors that reached 500 mm^3^ in size. One hour after injection, mice were euthanized, single-cell suspensions of tumors and tumor-draining lymph nodes (tdLN) were generated and were stained with cell surface markers followed by fixation and copper-catalyzed azide-alkyne cycloaddition of Alexa Fluor 647-azide (**Fig. 1c and Extended Data Fig. 1b**). Compared with vehicle-treated mice, single cells from OPP-treated mice showed a substantial increase in Alexa Fluor 647 fluorescence which was abrogated if mice were pre-treated with cycloheximide to block protein synthesis prior to OPP injection (**Fig. 1d**). OPP incorporation was significantly reduced in tumor-infiltrating CD44^+^CD8^+^ T-cells from E0771 tumor-bearing mice compared with either CD45^-^ cells within tumors or CD44^+^CD8^+^ T-cells within tumor-draining lymph nodes from the same mouse (**Fig. 1e**).

Intratumoral T-cell dysfunction is characterized by a conserved transcriptional program known as T-cell exhaustion, which develops over several weeks of tumor residence and is marked by sequential upregulation of inhibitory immunoreceptors including Programmed Cell Death-1 (PD-1) and T-cell immunoglobulin and mucin domain 3 (Tim-3) ^15–19^. Protein synthesis rates were decreased in CD8^+^ TILs regardless of inhibitory receptor expression, raising the possibility that suppression of TIL effector function might be independent of the T-cell exhaustion program (**Fig. 1f**). To test this hypothesis, we measured protein synthesis rates in activated OT-I^+^CD8^+^ T-cells from E0771-OVA tumors and tumor-draining lymph nodes (tdLN) 24 hours post adoptive cell transfer (**Fig. 1g**). CD8^+^ TILs exhibited reduced protein synthesis rates compared with CD8^+^ T-cells from the tdLN within 24 hours of adoptive transfer, suggesting that protein synthesis is suppressed in TILs prior to the development of canonical T-cell exhaustion (**Fig. 1h,i**).

Tumor-infiltrating CD8^+^ T-cells undergo persistent antigen exposure within a unique tumor microenvironment^20,21^. To test whether loss of protein synthesis in TILs requires both chronic TCR stimulation and exposure to the tumor microenvironment, we leveraged a widely used viral infection model in which mice are infected with variants of lymphocytic choriomeningitis virus (LCMV) that are either rapidly cleared (LCMV-Armstrong) or persistent (LCMV-Clone 13)^12^. T-cells from LCMV-Cl13 mice become exhausted over several weeks due to chronic antigen exposure. These cells can be found in secondary lymphoid organs including the spleen, allowing measurement of protein synthesis rates in exhausted T-cells in the absence of nutrient limitation. Antigen-specific CD8^+^ T-cells from the spleen of mice at 7 and 30 days following infection with LCMV-Cl13 show similar protein synthesis rates compared with T-cells from mice infected with the self-limited LCMV-Armstrong clone, despite increased expression of PD-1 and Tim-3 (**Extended Data Fig. 1c,d**). Taken together, these results indicate that tumor-specific suppression of CD8^+^ T-cell protein synthesis occurs rapidly upon tumor infiltration and is in large part environmentally driven.

### Intratumoral glutamine restriction limits CD8^+^ T-cell protein synthesis and cytotoxicity

We next asked how the tumor microenvironment suppresses TIL protein synthesis. In eukaryotic cells, protein synthesis requires transcription, splicing and export of messenger RNA (mRNA), assembly of ribosome and translation initiation complexes, transfer RNAs (tRNA), and sufficient amino acids and ATP/GTP to enable charging and transfer of amino acids to elongating peptide chains (**Extended Data Fig. 2a**). Passaging of activated T-cells in the presence of persistent T-cell receptor (TCR) stimulation, which has been shown to activate a transcriptional program similar to that seen in TILs^22–24^ did not compromise expression of cytotoxic mRNA transcripts such as *Ifng* or *Gzmb* (**Extended Data Fig. 2b**), translation initiation factors, ribosomal protein subunits, or transcript levels of cytosolic tRNA synthetases (**Extended Data Fig. 2c,d**)^22^. Similarly, expression of genes encoding translation initiation factors, ribosomal protein subunits, and cytosolic tRNA synthetases were not reduced in TILs compared with T cells from LCMV-Arm infected mice and were in fact generally increased in expression compared with T cells from LCMV-Cl13 infected mice (**Extended Data Fig. 2e**). In contrast, TILs isolated from either B16-F10 melanoma or E0771 mammary tumors upregulated genes associated with amino acid deprivation compared with T cells from LCMV-Arm infected mice (**Fig. 2a**). This signature was specific to tumor-infiltrating T-cells, as exhausted T-cells from LCMV-Cl13 infected mice did not upregulate genes associated with amino acid deprivation, suggesting that intratumoral amino acid availability might limit TIL protein synthesis. Consistent with this hypothesis, reducing extracellular amino acid availability was sufficient to limit both protein synthesis in T-cells encountering persistent antigen *in vitro* (**Fig. 2b,c**). To test whether decreased amino acid availability is sufficient to limit the cytotoxic capacity of T-cells during persistent antigen encounter, we tested the capacity of chronically stimulated T-cells to both release effector cytokines and kill cognate peptide-pulsed tumor cells *in vitro* (**Fig. 2d**). Amino acid limitation was sufficient to impair inflammatory cytokine release (**Fig. 2e**) as well as cytotoxicity (**Fig. 2f**), suggesting that amino acid limitation is sufficient to compromise intratumoral T-cell effector function and nominating amino acid insufficiency as a potential driver of rapid loss of TIL effector function *in vivo*.

**Fig 2.**
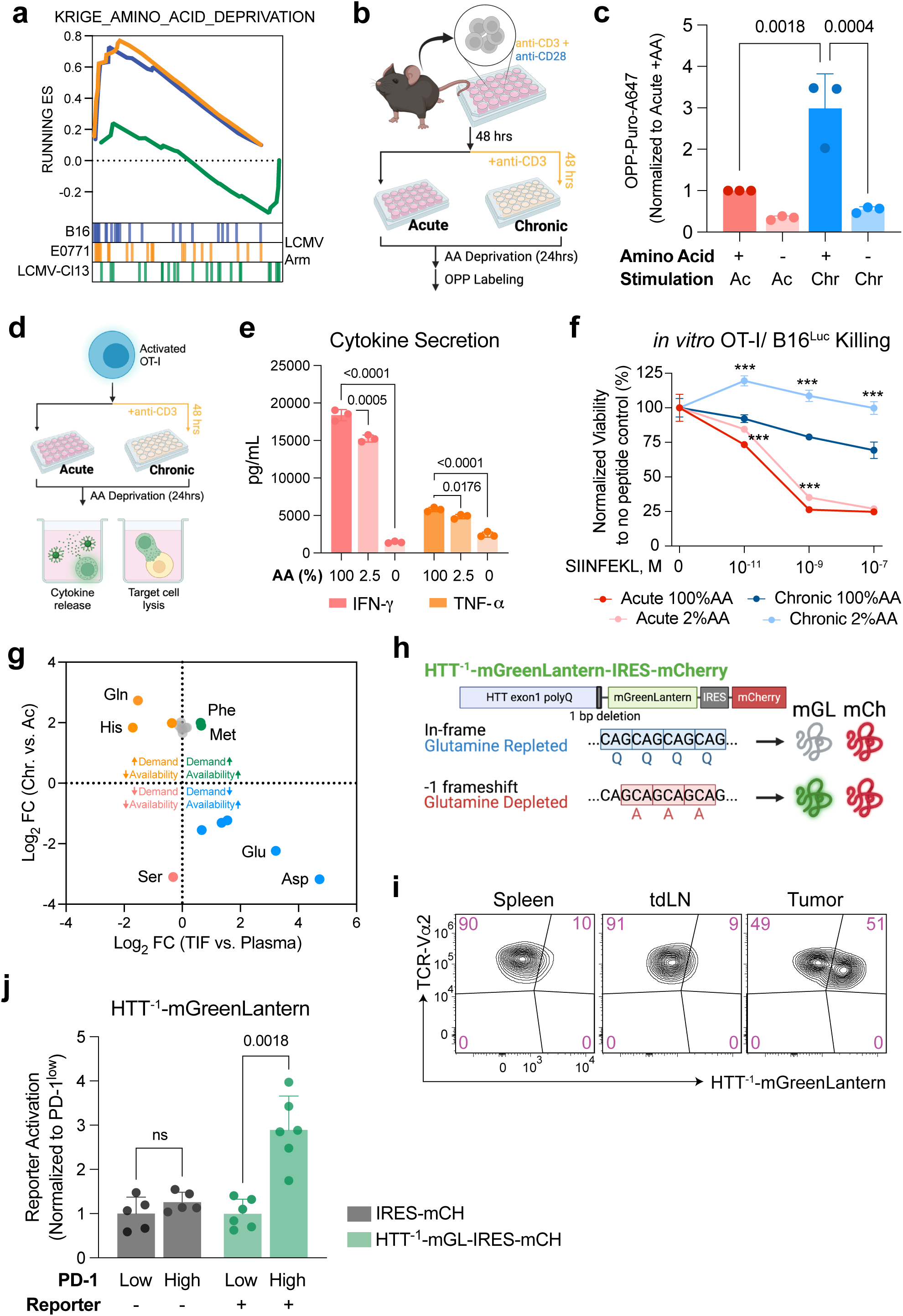
Intratumoral CD8^+^ T-cell protein synthesis is limited by glutamine availability. a, Gene set enrichment plot of KRIGE_AMINO_ACID_DEPRIVATION in CD8^+^CD44^+^PD-1^+^ T-cells from B16-F10 melanomas, E0771 breast tumors, or LCMV-Cl13 infected mice compared with CD8^+^CD44^+^ T-cells from LCMV-Armstrong infected mice. **b,** Experimental design to measure the impact of chronic TCR stimulation and amino acid deprivation on T-cell protein synthesis. **c,** Protein synthesis rates in acutely or chronically stimulated CD8^+^ T-cells cultured in media containing or lacking amino acids for 24 hours as indicated. **d,** Experimental design to assess cytokine release and cytotoxic capacity of amino acid-deprived T-cells *in vitro*. **e,** IFN-γ and TNF-α secretion of chronically stimulated T-cells cultured in the indicated amino acid conditions for 24 hours following re-stimulation with PMA and ionomycin. **f,** *In vitro* killing of B16-Luc cells pulsed with indicated concentration of SIINFEKL by acutely or chronically stimulated T-cells cultured in 100% or 2% amino acids for 24 hours. **g,** Comparison of chronic TCR stimulation-induced amino acid consumption (shown on y-axis with positive values reflecting increased consumption by chronically vs. acutely stimulated T-cells) with environmental amino acid availability (shown on x-axis with positive values reflecting increased abundance in TIF vs. plasma). Amino acid consumption is measured by subtracting metabolite abundances in cell culture supernatant from a media only control and normalizing by cell number in each sample. **h,** Diagram depicting an optimized reporter to detect glutamine insufficiency *in vivo* (HTT^-1^-mGreenLantern-IRES-mCherry). **i,** HTT^-1^-mGL fluorescence intensity of adoptively transferred TCR-Vα2^+^mCherry^+^ OT-I T-cells in spleen, tdLN, and tumor from B16-OVA tumor-bearing mice. **j,** mGreenLantern fluorescence in intratumoral TCR-Vα2^+^mCherry^+^ OT-I T-cells, normalized to PD-1-low T-cells. P values were calculated by one-way ANOVA with Tukey’s multiple comparisons test (**c**), Dunnett’s multiple comparisons test (**e**), or two-tailed Student’s t-test within groups (**f,j**). Data are presented as the mean ± s.d. of n = 3 biologically independent samples or n = 5 or 6 (**i,j**) independent mice from a representative experiment.

We next asked which amino acids were most likely to be limiting for T-cell function *in vivo*. We hypothesized that the functional sufficiency of any individual amino acid would be influenced by both the extracellular abundance and the intracellular consumption of that amino acid. To determine which amino acids were limited in extracellular availability, we measured the abundance of each amino acid within tumor interstitial fluid (TIF) relative to plasma from the same tumor-bearing mouse (**Extended Data Fig. 2f**). TIF samples from both B16 and E0771 tumors exhibited similar changes in metabolite composition relative to paired plasma samples, including established changes such as reduced glucose and increased lactate abundance (**Extended Data Fig. 2g-i**). Among amino acids, only glutamine, histidine, and serine were found to be substantially depleted within TIF **(Fig. 2g and Extended Data Fig. 2j)**. To determine which amino acids were most consumed by T-cells during persistent antigen encounter, we expanded activated T-cells in the presence of IL-2 alone or IL-2 plus persistent antigen stimulation and measured the depletion of amino acids from the culture media. Amongst all amino acids, we found that glutamine consumption was most strongly increased in T-cells expanded in the presence of persistent TCR stimulation compared with cytokine alone (**Fig. 2g and Extended Data Fig. 2k,l**), consistent with prior reports demonstrating increased glutamine uptake by TILs *in vivo* ^25^. Given the depletion of glutamine in TIF, these data suggested that TILs are functionally limited by glutamine availability within tumors.

To directly test whether intratumoral T-cells experience glutamine deprivation *in vivo*, we leveraged a recently published ribosomal frameshift-based fluorescent reporter of glutamine availability, which takes advantage of the fact that glutamine deprivation reduces translational fidelity of polyglutamine tract-containing proteins such as the Huntingtin protein (HTT)^26^. By introducing a single base deletion at the C-terminus of the HTT polyglutamine tract prior to a fluorescent reporter, fluorescent protein will only be translated if-1 ribosomal frameshifting occurs in response to glutamine deprivation^26^. We optimized this reporter by replacing GFP with a brighter, monomeric variant, mGreenLantern (mGL)^27^, and adding a frameshift-independent fluorescent reporter (mCherry) (**Fig. 2h**). Activated OT-I T-cells expressing this reporter (HTT^-1^-mGreenLantern-IRES-mCherry) were adoptively transferred into B16-OVA tumor-bearing Rag1KO recipient mice. 10 days post-adoptive transfer, over half (51.4%) of tumor-infiltrating, mCherry^+^ OT-I T-cells expressed mGL as compared with only 10% of mCherry^+^ OT-I T-cells in the spleen or draining lymph nodes (**Fig. 2i**). This resulted in a significant increase in mGL mean fluorescence intensity in tumor-infiltrating mCherry^+^ OT-I T-cells compared with mCherry^+^ OT-I T-cells from the dLN or spleen that was most substantial in PD-1^+^ TILs (**Fig. 2j and Extended Data Fig. 2m**). In contrast, no significant increase in mGL fluorescence was observed in activated splenic OT-I T-cells following adoptive transfer into mice infected with an ovalbumin-expressing strain of *Listeria monocytogenes* (Lm-OVA) (**Extended Data Fig. 2n**), confirming that T-cells experience glutamine limitation only within tumors. Moreover and in contrast to TILs, E0771 tumor cells stably transduced with HTT^-1^-mGreenLantern-IRES-mCherry did not express mGL *in vivo* despite robust induction upon amino acid limitation *in vitro* (**Extended Data Fig. 2o,p**). This lack of mGL induction was not due to vector-driven graft rejection as non-frameshift (HTT-mGL) expressing E0771 cells robustly activated mGL *in vivo* (**Extended Data Fig. 2p**). Thus, CD8^+^ TILs but not tumor cells become glutamine-limited within the tumor microenvironment. To identify the conditions required to drive glutamine limitation in T-cells upon tumor infiltration, we tested the impact of persistent antigen and amino acid deprivation, either alone or in combination, on HTT^-1^-mGreenLantern-IRES-mCherry-transduced T-cells *in vitro* (**Extended Data Fig. 2q**). We found that a combination of chronic TCR stimulation and amino acid deprivation were required for maximal mGL induction (**Extended Data Fig. 2r,s**). Thus, T-cell-specific glutamine insufficiency within tumors is caused by a combination of antigen-driven glutamine demand and TME-specific limitation of glutamine availability.

### Glutamine tRNA uncharging limits CD8^+^ T-cell function during amino acid limitation

We next asked how reduced glutamine availability suppresses intratumoral T-cell protein synthesis. Amino acid deprivation can suppress protein synthesis either through the uncharging of tRNA, leading to phosphorylation and activation of the general control nonderepressible 2 (GCN2) kinase-dependent stress response, or by preventing mammalian target of rapamycin complex 1 (mTORC1) activation at the lysosome (**Extended Data Fig. 3a**)^28,29^. Consistent with our finding that a combination of chronic TCR stimulation and amino acid withdrawal drives glutamine insufficiency and loss of protein synthesis (**Fig. 2c and Extended Data Fig. 2r,s**), we found that while 24 hours of amino acid withdrawal was sufficient to suppress mTORC1 activation in both acutely and chronically stimulated T cells, robust induction of GCN2 phosphorylation and downstream accumulation of activated transcription factor 4 (ATF4), a key mediator of the integrated stress response (ISR)^30^, required a combination of amino acid limitation and persistent antigenic stimulation (**Fig. 3a**). We therefore asked whether either activation of the ISR or inactivation of mTORC1 is required for amino acid-dependent suppression of protein synthesis. Treatment of T-cells with the ISR inhibitor ISRIB during chronic TCR stimulation increased protein synthesis rates when amino acids were freely available but did not restore protein synthesis when amino acids were limited, suggesting that amino acid-dependent suppression of protein synthesis is independent of the ISR (**Extended Data Fig. 3b**). Similarly, targeting mTORC1 inactivation via CRISPR-mediated editing of the GATOR1 component DEPDC5^31^ was sufficient to increase ribosomal S6 kinase (S6K1) phosphorylation when amino acids were present but did not restore protein synthesis during amino acid limitation (**Extended Data Fig. 3c,d**). Thus, amino acid limitation suppresses T-cell protein synthesis in a manner that is independent of mTORC1 silencing or ISR activation. We previously reported that tRNA^Gln^ uncharging under amino acid-limited conditions can independently suppress protein synthesis^26^. We therefore directly measured the impact of chronic TCR stimulation and amino acid availability on tRNA charging using a tRNA-isodecoder-specific quantitative PCR method that we previously developed^26^. Of the tRNA isodecoders tested, we observed the most substantial uncharging of tRNA^Gln^ in response to chronic TCR stimulation in a manner that was uniquely sensitive to acute amino acid withdrawal (**Fig. 3b**). Taken together, these findings indicate that glutamine limitation restricts CD8^+^ T-cell protein synthesis via tRNA uncharging.

**Fig 3.**
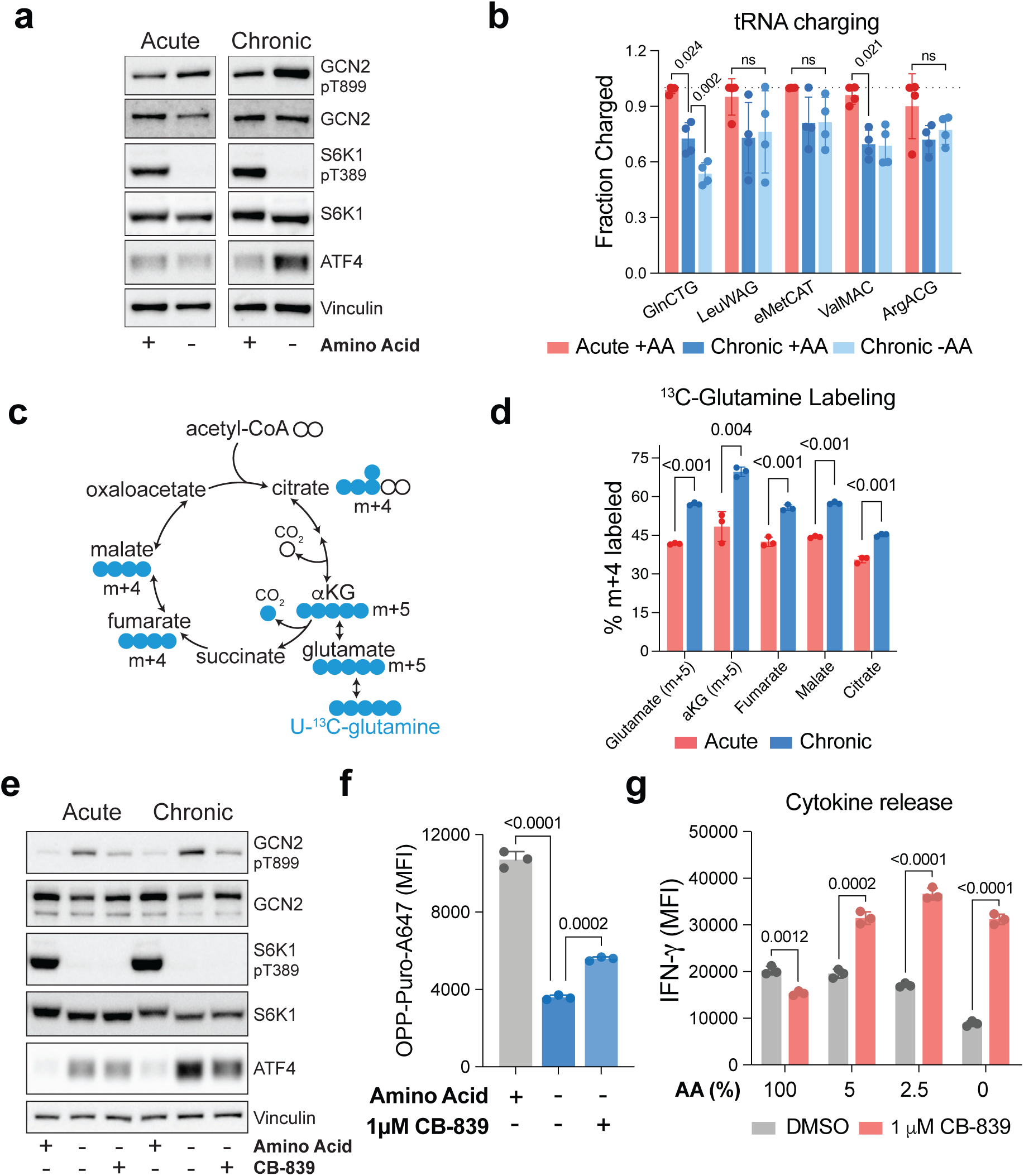
Glutamine tRNA uncharging limits CD8^+^ T-cell effector function. a, Amino acid stress response markers in acutely or chronically stimulated T-cells cultured in the presence or absence of amino acids for 6 hours, measured by Western blot. Vinculin was used as loading control. **b,** tRNA charging in acutely or chronically stimulated T-cells cultured in the presence or absence of amino acids for 6 hours using a tRNA isodecoder-specific qPCR method described in Methods. **c,** Schematic showing how oxidative metabolism of [U-^13^C]-Glutamine generates metabolites associated with the TCA cycle. Colored circles represent ^13^C-labeled carbons. **d,** Fractional labeling by [U-^13^C] glutamine of glutamate, α-ketoglutarate, fumarate, malate and citrate in acutely and chronically stimulated T cells. **e,** Amino acid stress response markers in acutely or chronically stimulated T-cells cultured in the presence or absence of amino for 6 hours, with or without the addition of the glutaminase inhibitor CB-839 measured by Western blot. Vinculin was used as loading control. **f,** OPP-AF647 MFI in chronically stimulated CD8^+^ T-cells cultured for 24h in the presence or absence of amino acids, with or without the addition of CB-839 (1µM). **g,** Cytokine release by chronically stimulated T-cells cultured in the indicated concentrations of extracellular amino acids for 24h, with or without the addition of CB-839. P values were calculated by one-way ANOVA with Tukey’s multiple comparisons test (**b,f**), Multiple two-tailed Student’s t-test (**d,g**). Data are presented as the mean ± s.d. of n = 3 biologically independent samples from a representative experiment, other than **a,e**, for which a representative result out of three independent experiments is shown.

### Glutaminase inhibition enhances T-cell function under amino acid limitation

Our observation that T-cells, but not cancer cells, experience glutamine limitation *in vivo* (**Fig. 2i and Extended Data Fig. 2p**) raised the question of why glutamine becomes uniquely limiting for chronically stimulated T-cells. To answer this question, we performed isotope tracing of stable, uniformly ^13^C-labeled glutamine ([U-^13^C]-Glutamine) in T-cells undergoing acute or chronic TCR stimulation *in vitro* (**Extended Data Fig. 3e**). Chronic TCR stimulation increased steady-state [U-^13^C]-Glutamine labeling of TCA cycle intermediates (**Fig. 3c,d**), leading us to hypothesize that glutamine anaplerosis in the TCA cycle might limit glutamine availability for tRNA charging and protein synthesis. To test this hypothesis, we treated T-cells with the glutaminase inhibitor CB-839, which limits glutamine anaplerosis by preventing glutamine deamidation to glutamate and has been shown to enhance anti-tumor immunity *in vivo*^32,33^. CB-839 treatment decreased GCN2 phosphorylation and ATF4 expression during amino acid limitation, demonstrating that glutamine anaplerosis is sufficient to drive tRNA uncharging and the amino acid stress response in chronically stimulated T-cells (**Fig. 3e**). As a result, CB-839 globally increased protein synthesis during amino acid limitation as measured by two independent methods (**Fig. 3f and Extended Data Fig. 3f**). Furthermore, CB-839 also decreased HTT^-1^-mGL reporter activation during amino acid limitation, indicating that glutaminase inhibition specifically improves intracellular glutamine availability in T-cells cultured in the absence of amino acids (**Extended Data Fig. 3g**). Accordingly, CB-839 was sufficient to restore intracellular IFN-γ production in chronically stimulated T-cells cultured under amino acid-limited conditions (**Fig. 3g**).

### Expression of a high-affinity amino acid transporter improves anti-tumor immunity *in vivo*

Glutamine supports intracellular amino acid homeostasis both directly and by serving as an exchange factor for several amino acid antiporters^34,35^ (**Fig. 4a**). We found that glutamine limitation was sufficient to decrease protein synthesis during chronic TCR stimulation (**Extended Data Fig. 4a**). However, supplementation of excess glutamine was insufficient to overcome global amino acid restriction (**Extended Data Fig. 4b**), suggesting that glutamine restriction suppresses protein synthesis in T-cells both by reducing tRNA^Gln^ charging as well as by limiting glutamine-dependent uptake of other amino acids. *SLC6A15* is a broad-spectrum neutral amino acid transporter that co-transports sodium with several essential amino acids (EAA), enabling cells to acquire EAA without exporting glutamine^36,37^. Accordingly, *SLC6A15* was found in a gain of function screen to enhance 293T cell proliferation during total amino acid restriction^38^. Overexpression of *SLC6A15* in OT-I T-cells, which do not express *SLC6A15* (**Extended Data Fig. 4c**) was sufficient to improve tumor control *in vivo* (**Fig. 4b,c**) and reduced markers of T-cell exhaustion during chronic stimulation *in vitro* (**Fig. 4d**). Mechanistically, *SLC6A15* overexpression had no impact on T-cell protein synthesis when amino acids were abundant, but restored protein synthesis when amino acid were limited (**Fig. 4e**). As a result, both cytokine release and killing of peptide-pulsed tumor cells were rescued by *SLC6A15* expression only when amino acids were limited (**Fig. 4f,g and Extended Data Fig. 4d**), indicating that *SLC6A15* enhances anti-tumor immunity by improving T-cell function during intratumoral amino acid restriction *in vivo*.

**Fig 4.**
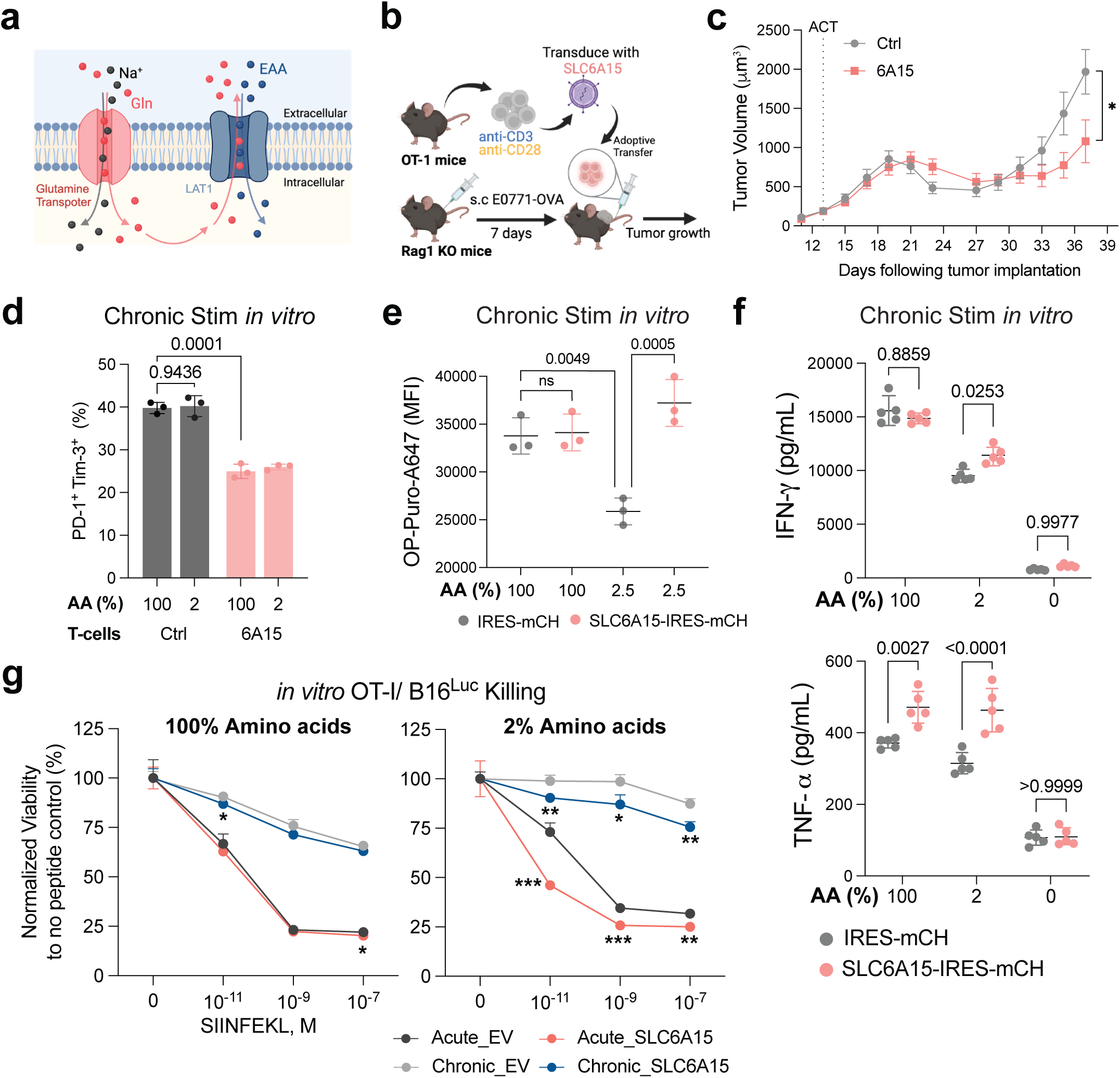
Expression of a high-affinity amino acid transporter improves anti-tumor immunity *in vivo*. a, Schematic depicting the role of glutamine in supporting amino acid homeostasis and transport. **b,** Experimental design to test whether *SLC6A15* overexpression in CD8^+^ T-cells improves anti-tumor immunity. **c,** Growth of subcutaneous E0771-OVA tumors following adoptive cell transfer (ACT) of OT-I T-cells expressing either vector control (Ctrl) or *SLC6A15* (6A15). **d,** Percentages of PD-1^+^Tim-3^+^ cells within CD44^+^CD8^+^ T-cell population expressing either empty vector (Ctrl) or *SLC6A15* (6A15) cultured in either 100% or 2% amino acid conditions in the presence of persistent TCR stimulation *in vitro*. **e,** OPP-AF647 MFI in Control or *SLC6A15*-overexpressing, chronically stimulated T-cells cultured in 100% or 2.5% amino acids for 24h. **f,** Cytokine release by Control or *SLC6A15*-overexpressing, chronically stimulated T-cells cultured in 100%, 2%, or 0% amino acids for 24 hours following re-stimulation with PMA and ionomycin. **g,** *In vitro* killing of B16-Luc cells pulsed with indicated concentrations of SIINFEKL by control or SLC6A15-overexpressing acutely or chronically stimulated T-cells cultured in 100% or 2% amino acids for 24 hours. P values were calculated by one-way ANOVA with Tukey’s multiple comparisons test (**d,e,f**), or multiple two-tailed Student’s t-test (**c,g**). Data are presented as the mean ± s.d. of n = 3 biologically independent samples or n = 7 independent mice from a representative experiment.

## Discussion

Loss of intratumoral cytotoxic function has been an established hallmark of cancer-associated T-cell dysfunction for over sixty years^8^. This dysfunction is accompanied by activation of a conserved transcriptional and epigenetic program known as T-cell exhaustion^6,8,9^. Whether the T-cell exhaustion program is required for loss of T-cell cytotoxic function remains unclear. Genetic deletion of the transcription factor TOX, which is required for the epigenetic and transcriptional remodeling of TILs, failed to restore cytokine production in TILs^3^. Moreover, TILs rapidly lose cytotoxic capacity despite adequate mRNA expression, suggesting independent, post-transcriptional regulators of cytotoxic T-cell function within the tumor microenvironment.

Here we establish a novel mechanism by which reduced amino acid availability within the tumor microenvironment can rapidly silence T-cell immunity. Loss of protein synthesis in TILs occurs within 24 hours of tumor infiltration, requires a combination of TCR engagement and amino acid limitation, and is driven primarily by glutamine-tRNA uncharging. This finding is consistent with recent reports showing both that glutamine is a major anaplerotic substrate for activated T cells *in vivo*^39^ and that intratumoral glutamine administration can restore anti-tumor immunity^40^. Whether the dependence of TILs on glutamine availability explain the poor response to immune checkpoint blockade in patients whose tumors express either high levels of Myc or a high abundance of cancer-associated fibroblasts, both of which drive glutamine depletion, is a subject of future exploration. Our findings also provide a mechanistic explanation for the long-standing observation that CD8^+^ TILs express high levels of transcripts encoding interferon gamma and granzymes but lack functional capacity. How amino acid availability regulates which mRNA are prioritized for translation is a question for future studies. Finally, the impact of blocking checkpoints such as PD-1 or LAG-3 on amino acid uptake within the tumor microenvironment is likely to inform the extent to which strategies to enhance intracellular amino acid availability in T-cells can synergize with immune checkpoint blockade to enhance immune-mediated tumor control.

## Acknowledgments

We thank members of the Vardhana lab for productive discussions that improved the manuscript. Y.C. was supported by an F99/K00 transition award (F99CA294267). SAV was supported by a Damon Runyon Clinical Investigator Award, a V Foundation Scholar Award, and a Mentored Career Development Award (NCI K08CA237731). We thank the Proteomics and Metabolomics Core Facility at Weill Cornell Medicine for assisting with metabolomics experiments.

## Author contributions

All experiments were designed by YC and SAV. YC performed all experiments with assistance from YL. JC assisted with metabolomics experiments. NP assisted with western blots, and tRNA charging assays. JR assisted with bioinformatic analyses. YC and SAV wrote the manuscript. All authors reviewed and approved of the manuscript prior to submission.

## Materials and correspondence

Correspondence and requests for materials should be addressed to SAV.

## Methods

### Cell lines and *in vitro* cultures

E0771, B16-F10, B16-Luc, B16-OVA were all purchased from ATCC. E0771-OVA cells were generated by introducing PresentER-SIINFEKL (mCherry), a gift from Dr. David Scheinberg (Addgene plasmid # 102945), into E0771 cells. E0771-LentiLucOS cells were generated by introducing pCDH-EF1a-eFFly-SIINFEKL-T2A-mCherry, cloned from Lenti-LucOS (Addgene plasmid # 22777, a gift from Dr. Tyler Jacks), into E0771 cells. All cell lines were cultured in DMEM + 10% FBS. All cells were routinely checked by the MycoAlert Mycoplasma Detection Kit (LT07-418) to be mycoplasma-negative.

### Plasmids

HTT^-1^-IRES-mGreenLantern (HTT^-1^-mGL) and HTT-IRES-mGreenLantern (HTT-mGL) were derived from the parental PolyQ^-1^-GFP ^26^ by replacing GFP with mGreenLantern, a brighter monomeric variant with longer half-life. In addition, the construct was subcloned into pMP71-IRES-mCherry-WPRE vector to enhance expression. pMSCV-SLC6A15-IRES-mCherry was generated by cloning human SLC6A15 cDNA into pMSCV-IRES-mCherry backbone. All vectors were amplified and purified from single colony and sequence-confirmed using Sanger sequencing.

### T-cell isolation and activation

Primary mouse T-cells were isolated from the spleen of C57BL/6J mice (Jackson Laboratory 000664) with the Dynabeads™ Untouched™ Mouse T Cells Kit (Invitrogen 11413D) following manufacturer’s instructions. Isolated T-cells were activated with plate-bound anti-CD3 antibody (2C11, 3 μg/mL, 3:1,000 dilution) and anti-CD28 antibody (37.51, 1 μg/mL, 1:1,000 dilution) for 2 days under RPMI-1640-based T-cell media (10% FBS, 2mM L-Glutamine, penicillin-streptomycin, 50µM 2-Mercaptoethanol, and 10 ng/mL of murine IL-2 (Peprotech)) and at a density of 1.5-2×10^6^ T-cell per mL.

### Acute and chronic T-cell stimulation

After 48 hours of T-cell activation described above, T-cells are replated in 6-well plates pre-coated with (chronic) or without (acute) anti-CD3 antibody (2C11, 3μg ml^−1^, 3:1,000 dilution) at a density of 1×10^6^ T-cell per ml in T-cell media (10% FBS, 2mM L-Glutamine, penicillin-streptomycin, 50µM 2-Mercaptoethanol, and 10 ng/mL of murine IL-2 (Peprotech)). For every 48 hours, cells are resuspended, spun down, and re-plated with fresh T-cell media with respective stimulating conditions. Chronically stimulated or time-matched acutely stimulated T-cells are harvested for experiments at the time stated in each individual experiment. Telaglenastat (CB-839, Selleck Chemicals LLC) was resuspended in dimethylsulfoxide (DMSO) and used at 1μM. O-Propagyl-Puromycin (OPP, Fisher Scientific) was resuspended in PBS and used at 10mM. ISRIB (MedChemExpress) was resuspended in dimethylsulfoxide (DMSO) and used at 0.4µM.

### Retroviral transduction

Plasmids expressing gene-of-interest were co-transfected with Moloney murine leukemia viral packaging plasmids (pCL-Eco) into Lenti-X 293T cell line (Takara) with the Lipofectamine transfection kit (Invitrogen) following the manufacturer’s instructions. Supernatants containing viral particles are collected 48 hours later, filtered through polyethersulfone (PES) filters, and precipitated overnight with 8% Polyethylene glycol (PEG) 8000 (Promega). Concentrated viral media is then added to 6-well plates pre-coated with RetroNectin reagent (Takara) (13.3μg/mL) and centrifuged to allow attachment of the viral particle onto the plate. Finally, primary mouse T-cells, activated as described in “T-cell isolation and activation” for 24 hours, are added to each well, followed by centrifugation at 300g for 10 min and incubated for 24 hours.

### CRISPR-mediated gene editing

Primary T-cells are isolated and activated from Rosa26^Cas9^ knock-in C57BL/6 mice (Jackson Laboratory 028555) as described in “T-cell isolation and activation”. After 24hrs, Cas9-expressing T-cells were transduced with retrovirus expressing empty vector control or 2 independent sgRNA targeting *Depdc5* (*sgDEPDC5#1*: GATTTAGTAAACAGGTCGGCG; *sgDEPDC5#2*: GCTCACCCCAATGATGAGTAC) following procedures described as above. Knock-out of gene of interest is confirmed through immunoblot analysis as well as flow cytometry.

### *In vitro* killing assay

B16 cells expressing Luciferase (B16-Luc) are first pulsed with SIINFEKL peptide (Biosynth) at indicated concentration for 2 hours at 37°C. Following one PBS wash, the peptide-pulsed B16-Luc cells are seeded in 96-well plates with 5×10^4^ cells per well. Acutely or chronically stimulated OT-I T-cells that are cultured under specified conditions are then added to each well at an effector:target ratio of 2:1 and incubated overnight at 37°C to enable killing. Cell Viability is measured the next day adding ONE-Glo™ Luciferase System Solution (Promega) to each well and measured with a GloMax 96 Microplate Luminometer (Promega). Cell viability is normalized to the no-peptide control for each condition.

### OPP-based translation measurement *in vitro*

For *in vitro* translation assessment, O-Propargyl-Puromycin (OPP) is spiked into cell culture at the final concentration of 10μM. After 30 minutes of incubation, cells are harvested and stained with TruStain FcX™ Fc block (Biolegend) and Ghost Dye™ Viability Dye (Tonbo), followed by staining with surface antibodies of interest. Cells are then fixed with pre-chilled methanol at-20°C for 10 minutes, washed with FACS buffer, and incubated with permeabilizing solution (0.5% Triton X in PBS) at room temperature for 15 minutes. Then, the cells are washed with FACS buffer and subjected to Click-iT™ Cell Reaction Buffer Kit (Invitrogen) and Andy Fluor™ 647 Azide (GeneCopoeia). Samples are washed again and analyzed immediately with Cytek^®^ Aurora (Cytek). Data analysis was performed using FlowJo v.10.9. All experiments were performed at least two independent times.

### Flow cytometry

For analysis of surface marker, cells are incubated with Fc Block (Biolegend) and Ghost Dye™ Viability Dye (Tonbo) for 10 minutes in phosphate-buffered saline (PBS). Following a wash with PBS containing 2% (w/v) FBS (FACS buffer), cells are stained with surface marker antibodies for 30 minutes. After an additional FACS buffer wash, cells are fixed and permeabilized with the FoxP3 Fixation/Permeabilization Kit (Thermo Fisher) for 30 minutes at room temperature, followed by incubation with intracellular antibodies overnight at 4°C. The next day, the cells were washed, fixed with 4% paraformaldehyde, and subjected to three additional washes before analysis on the Cytek^®^ Aurora (Cytek). For assessment of intracellular cytokine production, activated T-cells cultured under indicated conditions are re-stimulated with Cell Activation Cocktail (Biolegend). After 1 hour, Brefeldin A (Biolegend) is added to inhibit cytokine release. Three hours later, the cells were processed as described above for surface and intracellular marker staining before analysis on the Cytek^®^ Aurora (Cytek). Data analysis was performed using FlowJo v.10.9. All experiments were performed at least two independent times. Antibodies used are as follows-B220-BV496 (BD Biosciences, RA3-6B2, 612950), CD11b-BUV737 (BD Biosciences, M1/70, 612800), CD11b-BUV395 (BD Biosciences, M1/70, 563553), CD39-BUV395 (BD Biosciences, Y23-1185, 567264), CD39-PE-Dazzle 594 (Biolegend, Duha59, 143812), CD39-PE (Invitrogen, 24DMS1, 12-0391-80), CD4-BV711 (Biolegend, RM4-5, 100550), CD44-BB700 (BD Biosciences, IM7, 566506), CD44-FITC (eBiosiences, IM7, 11-0441-85), CD44-PerCP-Cy5.5 (Biolegend, IM7, 103032), CD45-BV570 (Biolegend, 30-F11, 103136), CD8-BV785 (Biolegend, 53-6.7, 100750), CD8-BUV615 (BD Biosciences, 53-6.7, 613004), F4/80-BV605 (Biolegend, BM8, 123133), Gzmb-AF700 (Biolegend, QA16A02, 372222), MHC-II-BV650 (Biolegend, M5/114.15.2, 107641), NK1.1-PE-Cy5 (Biolegend, PK136, 108716), NK1.1-BV750 (BD Biosciences, PK136, 746876), NK1.1-BUV615 (BD Biosciences, PK136, 751111), PD-1-APC-Cy7 (Biolegend, 29F.1A12, 135224), PD1-APC (Biolegend, RMP1-30, 109112), SLAMF6-BV605 (BD Biosciences, 13G3, 745250), TCF1-AF647 (Cell Signaling, C63D9, 6709S), TCRb-PerCP-Cy5.5 (Biolegend, H57-597, 109228), TCRβ-BV805 (BD Biosciences, H57-597, 748405), TCRvα2-PE-Cy7 (Biolegend, B20.1, 127822), Tim-3-BV421 (Biolegend, RMT3-23, 119723), TNF-BV650 (Biolegend, MP6-XT22, 506333), IFNγ-PE-Cy7 (Biolegend, XMG1.2, 505826). For measurement of cytokine release, T-cells cultured in indicated conditions are re-plated in respective fresh media and re-stimulated with Cell Activation Cocktail (Biolegend). After 48 hours, cells are spun down and measured for cell number, while the supernatant is diluted and subjected to BD™ Cytometric Bead Array kit (CBA) following the manufacturer’s instructions. The readout of the cytokine release is normalized to the cell number in each condition.

### Animal Model

All animal experiments were performed according to Memorial Sloan Kettering Cancer Center Institutional Animal Care and Use Committee guidelines (Protocol Number 20-10-012). For the analysis of polyclonal tumor-infiltrating T-cells, C57BL/6J (Jackson 000664) mice were implanted subcutaneously with 2 × 10^5^ E0771 cells. Tumor-bearing mice were monitored daily and euthanized when showing signs of morbidity. At the end-point of experiments, mice are euthanized with CO_2_ as described in the animal protocol, followed by cervical dislocation for confirmation of death. Tissues, such as tumors, spleen, tumor-draining lymph nodes, are dissected, minced, and digested with DNase I (Roche) and Liberase (Roche) in RPMI media at 37°C for 30 minutes. Digested tissues are meshed on a 100μm cell strainer, and the red blood cells are lysed in the Ammonium–chloride–potassium (ACK) lysing buffer. Single cell suspensions are then stained and analyzed with a flow cytometer as outlined in the “Flow Cytometry” section. For analysis of adoptively transferred transgenic antigen-specific T-cells, C57BL/6J Rag1 KO (Jackson 002216) mice were injected subcutaneously with 2 × 10^5^ B16-OVA or E0771-LentiLucOS cells depending on the experiments. 10 days post tumor-implantation, OT-I T-cells stimulated as outlined in “T-cell isolation and activation” were adoptively transferred to tumor-bearing mice through retro-orbital injection. For the SLC6A15 tumor control experiments, OT-I T-cells are transduced with SLC6A15 as mentioned in “Retroviral transduction” prior to the adoptive cell transfer. After adoptive transfer, tumor size is monitored and measured every other day. At end-point of the experiment, tissues are harvested as described above and analyzed on the flow cytometer. For analysis of cytokine production in tumor-infiltrating T cells *ex vivo,* B16-OVA cells were subcutaneously implanted into C57BL/6J Rag1 KO (Jackson 002216) mice. 10 days post-implantation, 1 × 10^6^ native OT-I transgenic T-cells are adoptively transferred to B16-OVA tumor-bearing mice. 10 days later, the mice were euthanized, and the tumors and tumor-draining lymph nodes were dissected and digested as described above. Following digestion, the tumor cell suspensions were resuspended in 40% Percoll (Cytiva) and centrifuged at 3,000rpm, with the break off, for 30 minutes. The cell debris floating in the supernatant was discarded and the live cell pellets were resuspended and treated with Ack buffer to remove the red blood cells. The resulting cells are counted and re-stimulated with Cell Activation Cocktail (Biolegend) for 1 hour, followed by the addition of Brefeldin A for another 3 hours as previously described. Cells are then process as outlined in the “Flow cytometry” section and analyzed on the Cytek^®^ Aurora (Cytek). Data analysis was performed using FlowJo v.10.9. All experiments were performed at least two independent times.

### *In vivo* cell type-specific translation measurement

To measure translation in endogenous polyclonal T-cells, 50mg/kg OPP or PBS vehicle control is injected intraperitoneally into E0771 tumor-bearing C57BL/6 mice. For cycloheximide (CHX) control, 30mg/kg CHX is injected into mice 15 minutes prior to OPP administration. After 1 hour of OPP incorporation, tissues of interest are dissected and enzymatically digested as outlined in “animal models” to prepare single cell suspension. Single cell suspensions are then processed as described in “OPP-based translation measurement *in vitro*” and analyzed with the Cytek^®^ Aurora (Cytek). For measuring translation rate in tumor infiltrating T-cells 24 hours post adoptive cell transfer, E0771-LentiLucOS cells are subcutaneously implanted in Rag1 KO mice. 21 days post-tumor implantation, 3 × 10^6^ OT-I transgenic T-cells activated as described in “T-cell isolation and activation” are adoptively transferred to tumor-bearing mice. After 24 hours, OPP is intraperitoneally injected to mice as described above and samples are prepared and analyzed with the Cytek^®^ Aurora (Cytek). Data analysis was performed using FlowJo v.10.9.

### Quantification of gene expression

RNA was extracted from T-cells with TRIzol (Invitrogen) and purified with Direct-zol RNA Microprep Kits (Zymo Research) according to the manufacturer’s instructions. 1μg of RNA was subjected to reverse transcription for cDNA using the iScript™ cDNA Synthesis Kit (Bio-Rad). Quantitative real-time PCR (qPCR) is performed using the Power SYBR Green PCR Master Mix (Applied Biosystems™) and SLC6A15-specific primers (fwd: GCATGTTCGGCACCATCGAG, rev: CGATGCCGTACACGAAGCAC; and fwd: CAGCATGGGTCATGGTGTGC, rev:GATCACGCCTCCGAATCCCA) with QuantStudio 7 Flex (Applied Biosystems). Actin was used as an endogenous control for all experiments and results were analyzed based on the ΔΔCt method. For quantification of gene expression *in vivo*, intratumoral CD8^+^CD44^+^ T-cells are sorted from B16 and E0771 tumor-bearing mice 21 days following tumor implantation. In addition, CD8^+^CD44^+^ T-cells are sorted from spleen of mice infected with LCMV Cl13 at 28 days post infection (dpi), and at 7 (dpi) with the self-limiting LCMV-Arm infected mice. RNA was then isolated from the sorted T-cells and processed for downstream analysis. RNA-seq samples were sequenced using the Illumina NextSeq 500 platform, generating 150-base pair paired-end reads. FASTQ files were quality-checked using FastQC v0.12.0 and trimmed with Trim Galore v0.6.10 to remove adapter sequences and read ends with Phred scores below 15. STAR v2.7.0a was used to align the FASTQ files to the mm10 reference genome. SAM files outputted by STAR were sorted using the sort function from Samtools v1.19.2, and indexed using the index function to generate BAM index files. A gene expression count matrix was generated using featureCounts v2.0.6, quantifying only read pairs that overlapped exons. Downstream differential gene expression analysis was performed using DESeq2 v1.42.0.

### Metabolomics Analyses

For metabolite extraction and targeted metabolite profiling, samples were extracted using pre-chilled 80% methanol (-80 °C). The extract was dried with a Speedvac, and redissolved in HPLC grade water before it was applied to the hydrophilic interaction chromatography LC-MS. Metabolites were measured on a Q Exactive Orbitrap mass spectrometer (Thermo Scientific), which was coupled to a Vanquish UPLC system (Thermo Scientific) via an Ion Max ion source with a HESI II probe (Thermo Scientific). A Sequant ZIC-pHILIC column (2.1 mm i.d. × 150 mm, particle size of 5 µm, Millipore Sigma) was used for separation of metabolites. A 2.1 × 20 mm guard column with the same packing material was used for protection of the analytical column. Flow rate was set at 150 μL/min. Buffers consisted of 100% acetonitrile for mobile phase A, and 0.1% NH_4_OH/20 mM CH_3_COONH_4_ in water for mobile phase B. The chromatographic gradient ran from 85% to 30% A in 20 min followed by a wash with 30% A and re-equilibration at 85% A. The Q Exactive was operated in full scan, polarity-switching mode with the following parameters: the spray voltage 3.0 kV, the heated capillary temperature 300 °C, the HESI probe temperature 350 °C, the sheath gas flow 40 units, the auxiliary gas flow 15 units. MS data acquisition was performed in the m/z range of 70–1,000, with 70,000 resolution (at 200 m/z). The AGC target was 1e6 and the maximum injection time was 250 ms. The MS data was processed using Xcalibur 4.1 (Thermo Scientific) to obtain the metabolite signal intensities. Identification required exact mass (within 5ppm) and standard retention times.

### Stable isotope tracing by ^13^C-glutamine

Acutely or chronically stimulated T cells were cultured as outlined in “Acute and chronic T cell stimulation” section. After 8 days, cells were washed with PBS and re-plated on plates coated with anti-CD3 antibody (3 μg/ml, 1:1000 dilution) in T-cell media. For isotopologue tracing studies, RPMI-1640 without glutamine was used, and supplemented with either ^12^C-glutamine (Gibco) or [U-^13^C] glutamine (Cambridge Isotope Laboratories) to a final concentration of 2 mM. After 6 hours, cells were harvested, centrifuged at 300g for 2 minutes at 4°C, and resuspended in 1 ml of ice-cold 80% methanol. After an overnight incubation at −80°C, samples were vortexed and centrifuged at 20,000g for 20 minutes to remove proteins. The supernatants were then dried using a vacuum evaporator (Genevac EZ-2 Elite) for 3 hours. The dried extracts were resuspended in either 60 μl of 60% acetonitrile in water for hydrophilic interaction liquid chromatography (HILIC) or 40 μl of 97:3 water/methanol containing 10 mM tributylamine and 15 mM acetic acid for ion pair liquid chromatography separations. Samples were vortexed, incubated on ice for 20 minutes, and then clarified by centrifugation at 20,000g for 20 minutes at 4°C. HILIC LC–MS analysis was carried out on a 6545 Q-TOF mass spectrometer (Agilent Technologies) in both positive and negative ionization modes, using previously described columns, buffers, and LC–MS parameters. Targeted data analysis, isotopologue extraction, and natural isotope abundance correction were performed with MassHunter Profinder software v.10.0 (Agilent Technologies). Ion pair LC–MS analysis was conducted with LC separation on a Zorbax RRHD Extend-C18 column (150 × 2.1 mm, 1.8-μm particle size, Agilent Technologies) and employed a gradient of solvent A (10 mM tributylamine and 15 mM acetic acid in 97:3 water/methanol) and solvent B (10 mM tributylamine and 15 mM acetic acid in methanol), following the manufacturer’s instructions (MassHunter Metabolomics dMRM Database and Method, Agilent Technologies).

### Amino acid consumption and availability measurement

For *in vitro* amino acid consumption assessment, T-cells are isolated and activated as described in “T-cell isolation and activation”. On day 2 following initial activation, cells are re-plated on plates coated with (chronic) or without (acute) anti-CD3 antibody (3 μg ml^−1^, 3:1,000 dilution) in T-cell media. In addition, media-only wells are included on the same plate and incubated without T-cells for the baseline of nutrient abundance. After 2 days, T-cell cultures are harvested and centrifuged at 300g for 5 min to collect the supernatant. Samples are processed and analyzed as outlined in “Metabolomic analyses”. For *in vivo* nutrient availability measurement in the tumor microenvironment, B16-OVA or E0771 cells are implanted subcutaneously into the flank of C57BL/6J mice. Mice were euthanized when tumor volume reached 1,000mm^3^ and blood was collected through cardiac puncture in EDTA collection tubes. Plasma was isolated after spinning at 845g for 10 minutes at 4°C and snap-frozen. To isolate tumor-interstitial fluid^41^, tumors were dissected after cardiac puncture, rinsed in ice cold saline, and rolled around on a filter paper to remove the excess saline. Tumors were then placed on top of a 20μm cell strainer and centrifuged at 106g for 10 minutes at 4°C. After spin, the supernatants were collected and snap-frozen in liquid nitrogen until the time of analysis. Samples are processed and analyzed as outlined in “Metabolomic analyses”.

### Immunoblotting

T-cells were harvested and lysed in RIPA buffer (Thermo) on ice and centrifuge at 13,000g for 20 minutes to collect clear cell lysate. Same amount of lysates across different samples are loaded, separated by SDS-PAGE, and transferred to nitrocellulose membranes (Bio-Rad). The membranes were blocked with 5% milk in Tris (pH)-buffered saline with 0.1% Tween-20 (TBST) and incubated overnight with primary antibodies at 4°C. The next day, membranes were washed three times with TBST and incubated with respective horseradish peroxidase (HRP)-conjugated secondary antibodies for 1 hour at room temperature on a shaker. After incubation with 1-shot digital-enhanced chemiluminescent (ECL) substrate (Kindle Bioscience), the membranes were imaged with a ChemiDoc Touch Imaging System (Bio-Rad). The antibodies used were: p-Thr899 GCN2 (Abcam, ab75836), GCN2 (Cell Signaling, 3302S), p-Thr389 S6K1 (Cell Signaling, 9234S), S6K1 (Cell Signaling, 2708S), ATF4 (Cell Signaling, 11815), Vinculin (Sigma, V9131), eIF4G (Thermo, PA5-17422), eIF4A1 (Thermo, 711505), eIF4E (Thermo, MA1-089), RPS3 (Cell Signaling, 2579S), RPL7a (Cell Signaling, 2415S), RPL26 (Cell Signaling, 2065S), β-Actin (Sigma, A2228), DEPDC5 (GeneTex, GTX133570), Puromycin (Sigma, MABE343).

### tRNA charging assay

Acutely or Chronically stimulated T-cells were cultured in indicated amino acid drop-out media for 6 hours, resuspended, washed with ice cold PBS, and lysed on ice with TRIzol (Invitrogen). After cell lysis, 0.2x volume of chloroform was added, shaken, centrifuged at 18,600g, and precipitated overnight using 2.7x volume of cold ethanol with 30 µg of GlycoBlue Coprecipitant (Thermo). The samples were resuspended in 0.3M acetate buffer (pH 4.5) containing 10 mM EDTA and precipitated again. The following day, the resulting sample were resuspended in 10 mM acetate buffer with 1 mM EDTA. 2 µg of each RNA sample was treated with 10 mM sodium periodate (Sigma-Aldrich) (“oxidized samples”) or with sodium chloride (“non-oxidized samples”) and incubated in the dark at room temperature for 20 minutes. The reactions were quenched with glucose for 15 minutes, and yeast tRNA^Phe^ (Sigma-Aldrich) was added prior to precipitating the samples with ethanol. The samples were then resuspended in 50 mM Tris buffer (pH 9) and incubated at 37°C for 50 minutes, followed by quenching with acetate buffer and another round of precipitation. Lastly, the samples were resuspended in RNase-free water and ligated to a 5’-adenylated DNA adaptor (5’-/5rApp/TGGAATTCTCGGGTGCCAAGG/3ddC /-3’) using truncated KQ mutant T4 RNA ligase 2 (New England Biolabs) for 3 hours at room temperature, following the protocol by Loayza-Puch et al., 2016. Reverse transcription was performed with a primer complementary to the DNA adaptor using SuperScript IV reverse transcriptase (Thermo) according to the manufacturer’s instructions. The cDNA samples were then analyzed by qPCR using tRNA isodecoder-specific primers, where the forward (FW) primer matched the 5’ end of the tRNA and the reverse (RV) primer spanned the junction between the 3’ end of the tRNA and the ligated adaptor. The following primer pairs were used:

ValMAC FW: 5’-GTTTCCGTAGTGTAGTGGTTATCACGTTCG-3’ RV: 5’-GAGAATTCCATGGTGTTTCCGCCC-3’

iMetCAT FW: 5’-AGCAGAGTGGCGCAGCG-3’

RV: 5’-GAGAATTCCATGGTAGCAGAGGATGGTTTCG-3’ eMetCAT FW: 5’-GCCTCSTTAGCGCAGTAGGTAG-3’

RV: 5’-GAGAATTCCATGGTGCCCCSTS-3’

GlnCTG FW: 5’-GGTTCCATGGTGTAATGGTNAGCACTCTG-3’ RV: 5’-GAGAATTCCATGGAGGTTCCACCGAGATTTG-3’

LeuWAG FW: 5’-GGTAGYGTGGCCGAGCG-3’

RV: 5’-GAGAATTCCATGGCAGYGGTGGG-3’

ArgACG FW: 5’-GGGCCAGTGGCGCAATG-3’

RV: 5’-GAGAATTCCATGGCGAGCCAGC-3’

Primers were designed based on reference tRNA sequences from the GtRNA database (http://gtrnadb.ucsc.edu/, RRID) (Chan and Lowe, 2016). Yeast tRNA^Phe^ were used as a control and the Ct values obtained using primers specific to yeast tRNA^Phe^ were subtracted from those obtained with primers targeting the tRNA isodecoder of interest. The charged fraction was determined by calculating the relative difference between the delta-Ct values from non-oxidized samples (representing total tRNA) and oxidized samples (representing charged tRNA) for each primer pair.

## Statistical analysis

GraphPad PRISM 10 software was used for statistical analyses. Error bars, P values and statistical tests are reported in figure legends.

