## Supplementary Figures for "Intratumoral amino acid insufficiency limits CD8^+^ T-cell effector function"

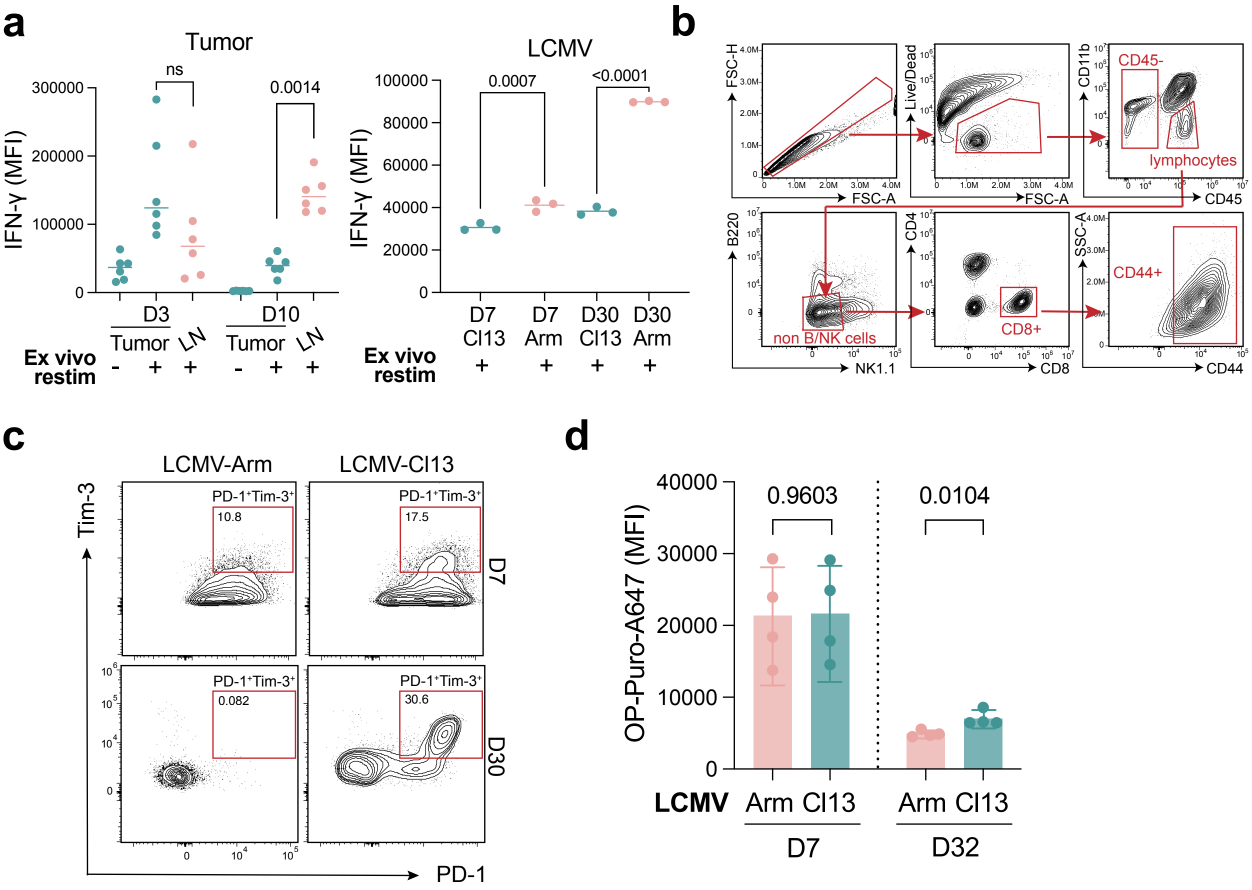


**Extended Data Figure 1. Measurement of T cell specific protein synthesis in vivo. a,** Left, IFN-γ expression in adoptively transferred CD8⁺CD44⁺ OT-I T-cells from tumors and tumor-draining lymph nodes at 3- or 10-days post adoptive transfer in the presence or absence of restimulation with PMA and Ionomycin. Right, IFN-γ expression in CD8⁺CD44⁺ T-cells from mice infected with either LCMV Armstrong or Clone 13 at 7- or 30-days post-infection upon restimulation with PMA and Ionomycin. **b,** Gating strategy for OPP-A647 measurements in CD8⁺CD44⁺ T-cells from tumors and tumor-draining lymph nodes. **c,** Expression of PD-1 and Tim-3 on CD8⁺CD44⁺ T-cells from mice at 7- or 32-days following infection with LCMV-Armstrong or LCMV-Clone13. **d,** OPP-A647 in CD8⁺CD44⁺ T-cells from mice at 7- or 32-days following infection with LCMV-Armstrong or LCMV-Clone13. P values were calculated two-tailed Student's t-test **(d)**. Data are presented as the mean ± s.d. of n = 6 **(a, left)**, 3 **(a, right)**, or 4 **(d)** independent mice from a representative experiment.

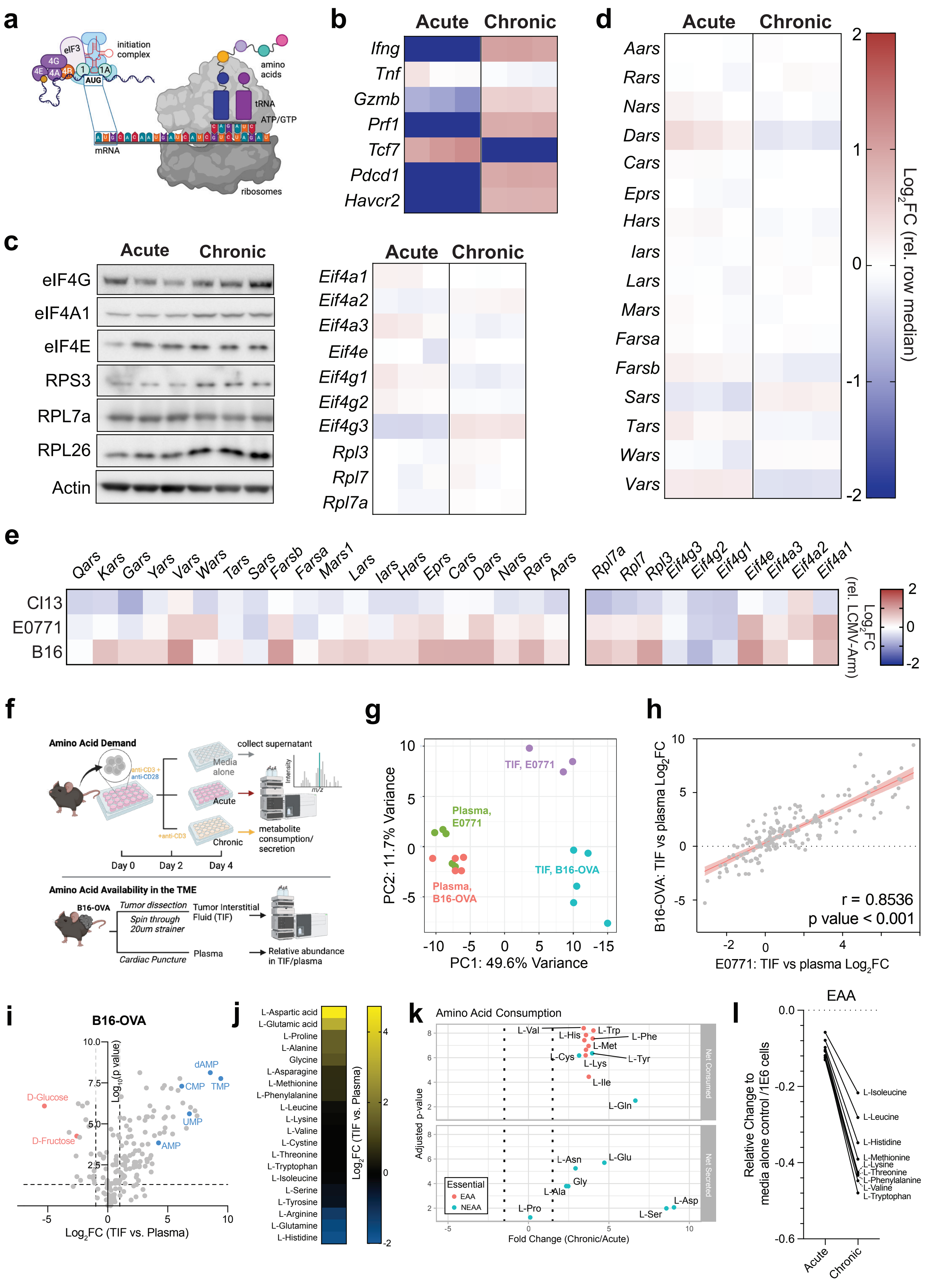
a
eIF3
4G
4E
4A
4B
1
1A
AUG
initiation complex
amino acids
tRNA
ATP/GTP
mRNA
ribosomes
b
Acute
Chronic
Ifng
Tnf
Gzmb
Prf1
Tcf7
Pdcd1
Havcr2
c
Acute
Chronic
eIF4G
eIF4A1
eIF4E
RPS3
RPL7a
RPL26
Actin
Acute
Chronic
Eif4a1
Eif4a2
Eif4a3
Eif4e
Eif4g1
Eif4g2
Eif4g3
Rpl3
Rpl7
Rpl7a
d
Acute
Chronic
Aars
Rars
Nars
Dars
Cars
Eprs
Hars
Iars
Lars
Mars
Farsa
Farsb
Sars
Tars
Wars
Vars
Log2FC (rel. row median)
2
1
0
-1
-2
e
Qars Kars Gars Yars Vars Wars Tars Sars Farsb Farsa Mars1 Lars Iars Hars Eprs Cars Dars Nars Rars Aars
Rpl7a Rpl7 Rpl3 Eif4g3 Eif4g2 Eif4g1 Eif4e Eif4a3 Eif4a2 Eif4a1
Cl13
E0771
B16
Log2FC (rel. LCMV-Arm)
2
0
-2
f
Amino Acid Demand
anti-CD3 + anti-CD28
collect supernatant
Media alone
Acute
+anti-CD3
Chronic
Intensity
m/z
metabolite consumption/ secretion
Day 0
Day 2
Day 4
Amino Acid Availability in the TME
B16-OVA
Tumor dissection
Spin through 20um strainer
Tumor Interstitial Fluid (TIF)
Cardiac Puncture
Plasma
Relative abundance in TIF/plasma
g
TIF, E0771
Plasma, E0771
TIF, B16-OVA
Plasma, B16-OVA
PC2: 11.7% Variance
PC1: 49.6% Variance
h
B16-OVA: TIF vs plasma Log2FC
E0771: TIF vs plasma Log2FC
r = 0.8536
p value < 0.001
i
B16-OVA
D-Glucose
D-Fructose
dAMP
CMP
TMP
UMP
AMP
Log10(p value)
Log2FC (TIF vs. Plasma)
j
L-Aspartic acid
L-Glutamic acid
L-Proline
L-Alanine
Glycine
L-Asparagine
L-Methionine
L-Phenylalanine
L-Leucine
L-Lysine
L-Valine
L-Cystine
L-Threonine
L-Tryptophan
L-Isoleucine
L-Serine
L-Tyrosine
L-Arginine
L-Glutamine
L-Histidine
Log2FC (TIF vs. Plasma)
k
Amino Acid Consumption
L-Val
L-His
L-Trp
L-Phe
L-Met
L-Cys
L-Lys
L-Tyr
L-Ile
L-Gln
Net Consumed
L-Asn
L-Glu
Gly
L-Ala
L-Pro
L-Asp
L-Ser
Net Secreted
Essential
EAA
NEAA
Adjusted p-value
Fold Change (Chronic/Acute)
l
EAA
Relative Change to media alone control /1E6 cells
L-Isoleucine
L-Leucine
L-Histidine
L-Methionine
L-Lysine
L-Threonine
L-Phenylalanine
L-Valine
L-Tryptophan
Acute
Chronic

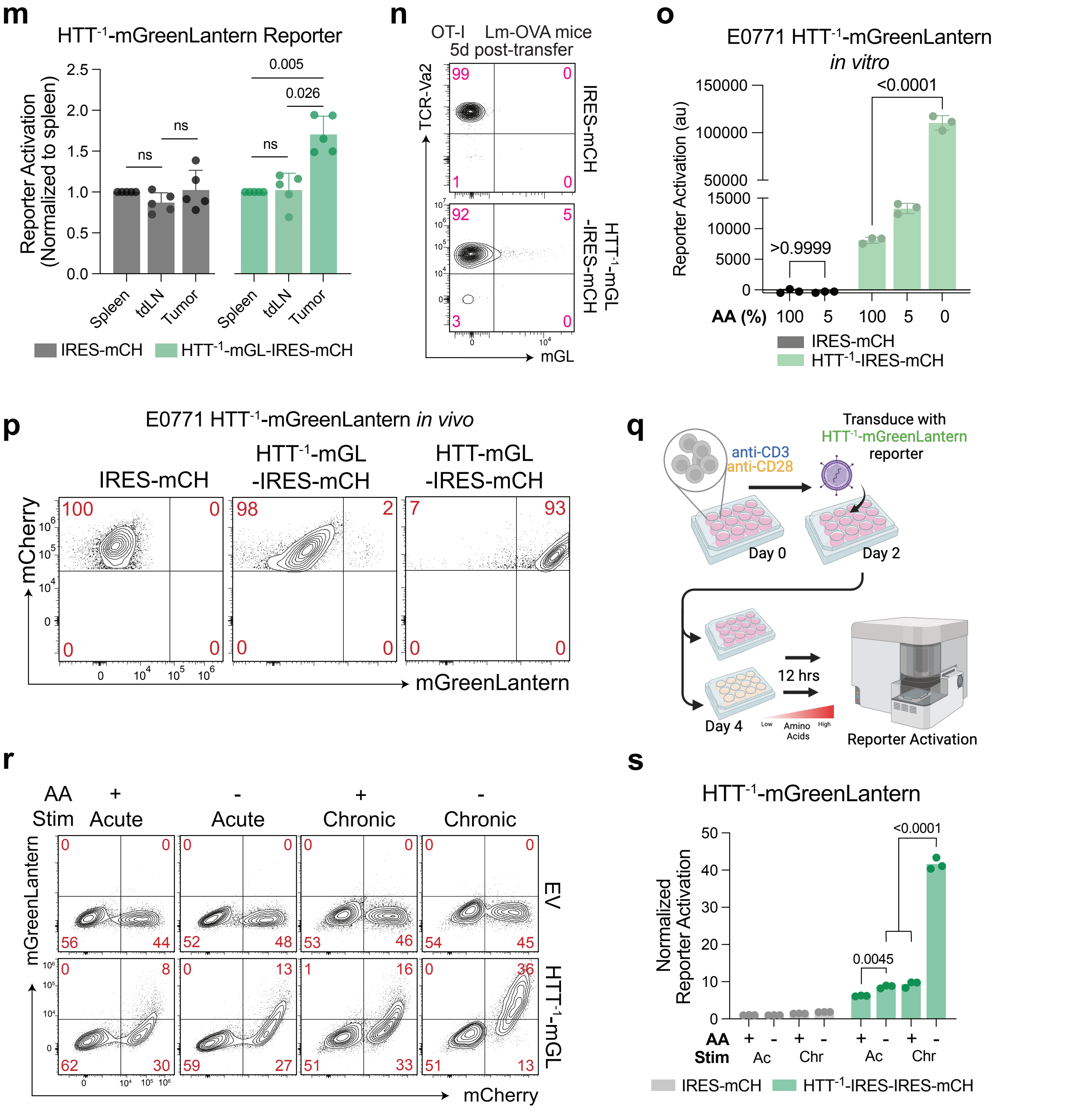


**Extended Data Figure 2. Tumor-infiltrating T-cells are amino acid limited for protein synthesis in vivo. a,** Schematic depicting mRNA translation with key components indicated. **b,** Heatmap showing the expression of T cell exhaustion-associated genes in activated primary mouse T-cells following 6 days of culture in the presence of cytokine alone (acute) or cytokine + anti-CD3 (chronic) measured by RNA-seq. **c,** Left, Western blot showing expression of cap initiation and ribosomal proteins in acutely or chronically stimulated T-cells. Actin was used as loading control. A representative result out of three independent experiments is shown. Right, heatmap showing expression of cap initiation and ribosomal proteins in acutely or chronically stimulated T-cells measured by RNA-seq. **d,** Heatmap showing expression of cytosolic tRNA synthetases in acutely or chronically stimulated T-cells measured by RNA-seq. **e,** Heatmap showing the expression of cytosolic tRNA synthetases, cap initiation and ribosomal proteins in $CD8^+CD44^+PD\text{-}1^+$ T-cells from B16-F10 melanomas, E0771 breast tumors, or LCMV-Cl13 infected mice compared with $CD8^+CD44^+$ T-cells from LCMV-Armstrong infected mice. **f,** Experimental design for measuring amino acid consumption during chronic stimulation and amino acid availability within tumors. **g,** Principal component analysis of steady state metabolite abundances in TIF and plasma extracted from E0771 or B16-OVA tumor-bearing mice. **h,** Pearson correlation coefficient of metabolite enrichment in TIF versus plasma in E0771 or B16-OVA tumor-bearing mice. **i,** Quantification of relative metabolite abundances in TIF compared with plasma in B16-OVA tumor-bearing mice. Colored circles indicate selected metabolites (red: carbohydrates; blue: nucleotides). **j,** Heatmap displaying relative abundance of amino acids in TIF versus plasma from B16-OVA tumor-bearing mice. Color scale represents $Log_2$ fold change relative to plasma. **k,** Quantification of relative metabolite abundances in media after culturing acutely or chronically stimulated T-cells. Fold change is calculated by subtracting relative metabolite abundances in each media with the metabolite abundances in the media alone control as described in Methods. **l,** Consumption of essential amino acids (EAA) in acutely or chronically stimulated T-cells per 1E6 cells. **m,** mGreenLantern fluorescence in $TCR\text{-}Va2^+mCherry^+$ OT-I T-cells within the draining LN and tumor, normalized to fluorescence in cells isolated from the spleen for each vector (iRES-mCharry and $HTT^{-1}$-mGL-iRES-mCherry), respectively. **n,** Fluorescence intensity of mGreenLantern in $HTT^{-1}$-mGL-iRES-$mCherry^+$ TC`R-$V\alpha 2^+$ T-cells 6 days after adoptive transfer into Lm-OVA infected-mice. **o,** Quantification of iRES-mCherry or $HTT^{-1}$-mGL-iRES-mCherry transduced E0771 cells cultured in media containing the indicated concentrations of amino acids for 24 hours. **p,** Fluorescence intensity of E0771 cells expressing IRES-mCherry control, $HTT^{-1}$-mGL IRES-mCherry, or a non-frameshifted variant, HTT-mGL iRES-mCherry following subcutaneous implantation in the flank of C57/Bl6 mice. Tumors were isolated when they reached 500 $mm^3$ in size. **q,** Schematic of experimental design for testing $HTT^{-1}$-mGL reporter activation in acutely or chronically stimulated T-cells cultured with or without extracellular amino acids. **r,s,** Reporter expression **(r)** and quantification **(s)** of iRES-mCherry or $HTT^{-1}$-mGL-iRES-mCherry transduced acutely or chronically stimulated T-cells cultured with or without extracellular amino acids for 24 hours. P values were calculated by one-way ANOVA with Tukey's multiple comparisons **(m,o,s)**. Data are presented as the mean ± s.d. of n = 3 biologically independent samples or n >3 independent mice from a representative experiment. Color scale for **(b,c,d,e)** represents log2 fold change relative to row median.

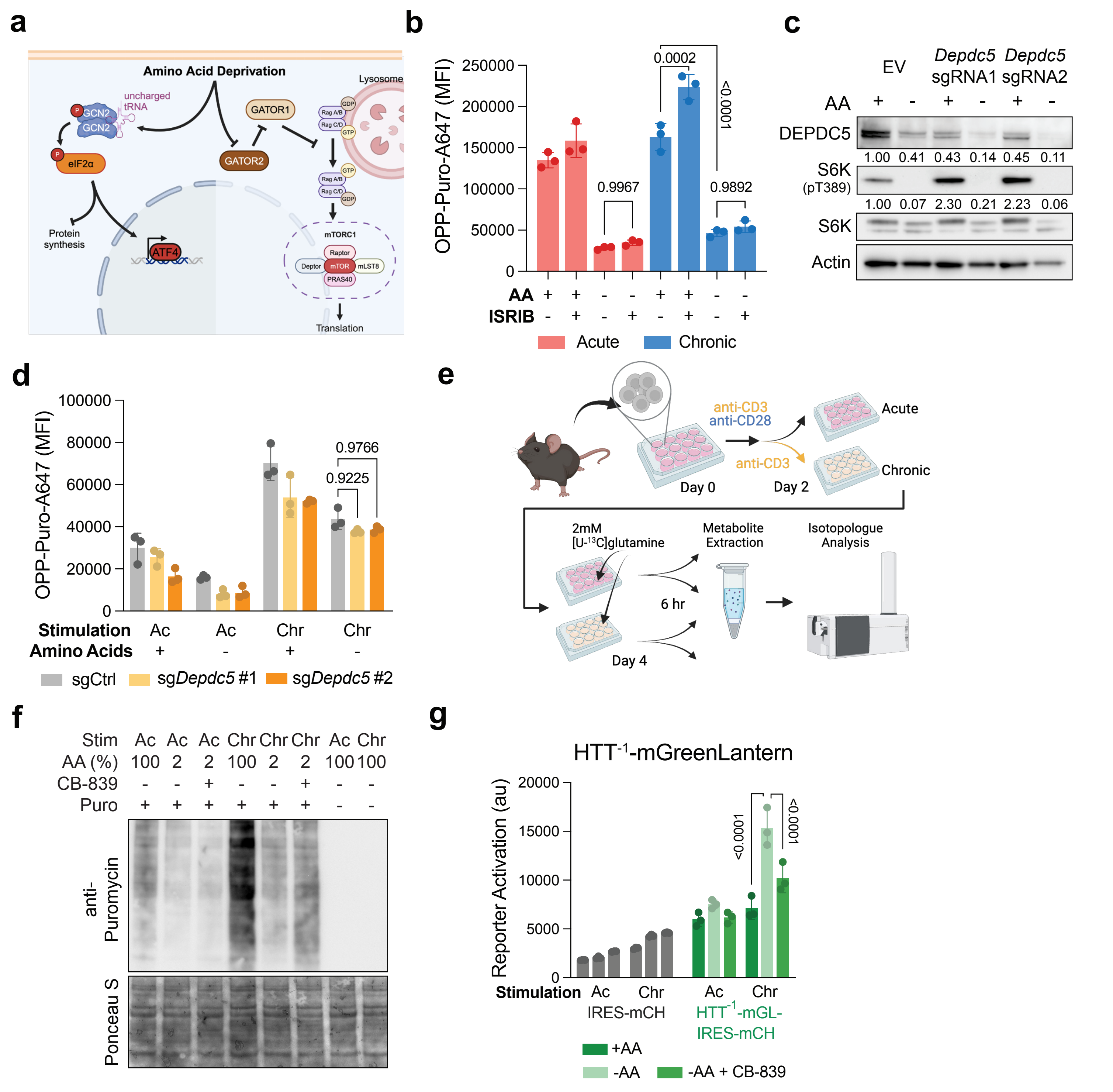

**Extended Data Figure 3. tRNA uncharging suppresses T-cell protein synthesis during amino acid limitation. a,** Schematic depicting two primary pathways governing amino acid regulation of protein synthesis. **b,** OPP-AF647 MFI in acutely or chronically stimulated CD8$^{+}$ T-cells cultured for 24 hours in the presence or absence of extracellular amino acids with or without the addition of 0.4μM ISRIB. **c,** Western blot of DEPDC5, and S6K1 phosphorylation in control or DEPDC5 KO T-cells cultured in the presence or absence of extracellular amino acids. Actin was used as loading control. A representative result out of three independent experiments is shown. Value below each blot represents the normalized abundance of each protein compared with control cells cultured in the presence of amino acids. **d,** OPP-AF647 MFI in acutely or chronically stimulated Cas9-expressing CD8$^{+}$ T-cells with or without CRISPR-mediated Depdc5 KO cultured in the presence or absence of extracellular amino acids for 24 hours. **e,** Experimental design to investigate changes in glutamine utilization during the presence of persistent TCR stimualtion in T-cells. **f,** Western blot showing puromycin incorporation into newly translated proteins in acutely or chronically stimulated T-cells cultured in media containing or lacking amino acids for 24 hours in the presence or absence of 1μM CB-839. Ponceau S staining on the same blot was used as loading control. **g,** mGreenLantern expression in mCherry$^{+}$ acutely or chronically stimulated T-cells transduced with vectors containing iRES-mCherry alone or HTT$^{-1}$-mGL iRES-mCherry, cultured for 24 hours in media containing or lacking amino acids in the presence or absence of 1μM CB-839. P values were calculated by one-way ANOVA with Tukey's multiple comparisons test **(b,d,g)**. Data are presented as the mean ± s.d. of n = 3 biologically independent samples from a representative experiment.

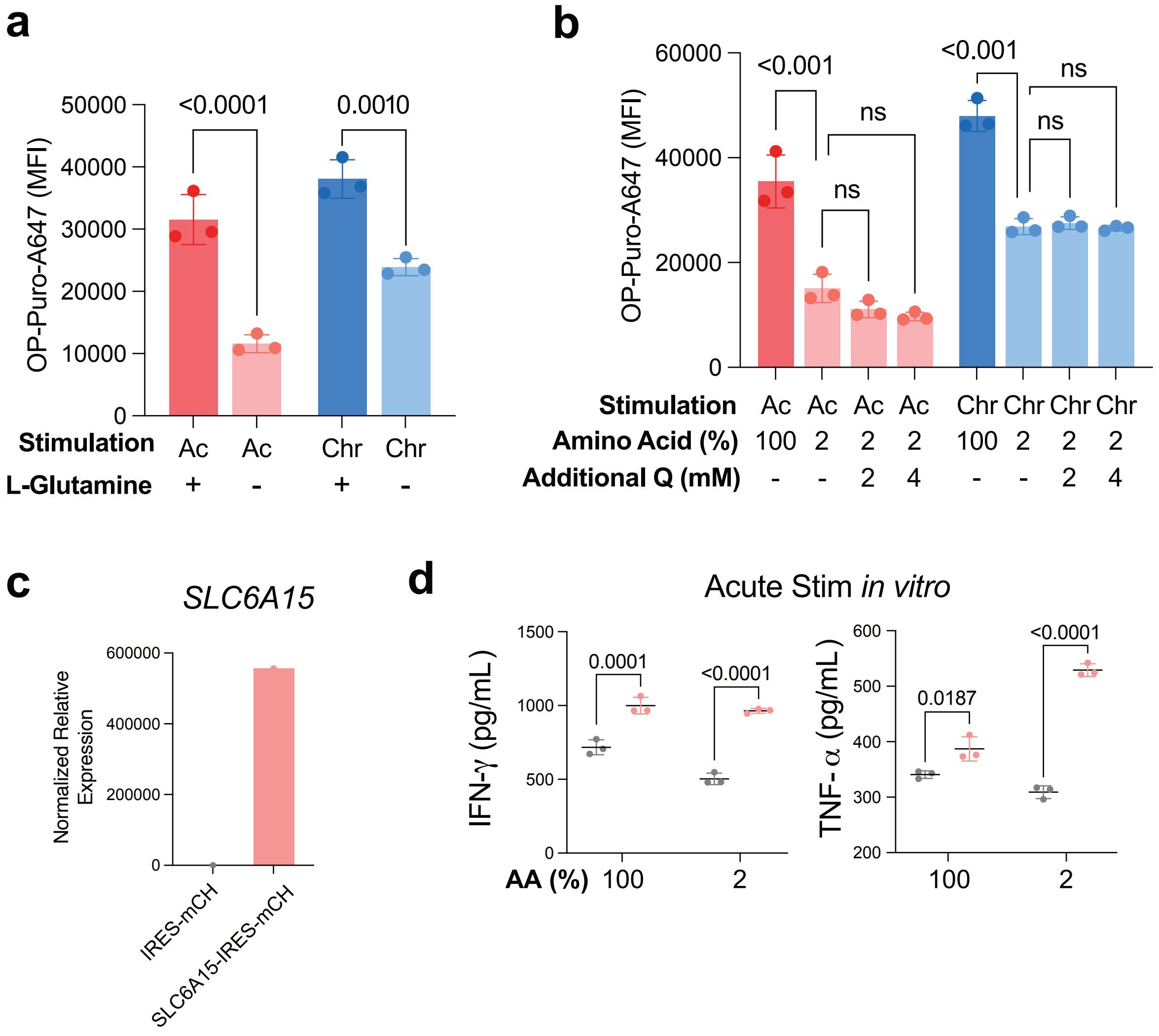


**Extended Data Fig. 4. Impact of SLC6A15 overexpression in primary T-cells. a,** Protein synthesis rates in acutely or chronically stimulated CD8$^+$ T-cells cultured in media containing or lacking glutamine for 24 hours as indicated. **b,** Protein synthesis rates in acutely or chronically stimulated CD8$^+$ T-cells cultured in media containing complete or 2% amino acids supplemented with either 2mM or 4mM of glutamine. **c,** qRT-PCR of SLC6A15 in T-cells transduced with either vector control or pMSCV-SLC6A15-IRES-mCherry. **d,** Cytokine release by Control or SLC6A15-overexpressing, acutely stimulated T-cells cultured in 100% or 2% amino acids for 24 hours following re-stimulation with PMA and ionomycin. P values were calculated by Tukey's multiple comparisons test **(a,b,c)**. Data are presented as the mean ± s.d. of n = 3 biologically independent samples or n = 7 independent mice from a representative experiment.
